# Oligodendrocyte precursor cells engulf synapses in a model of the developing human forebrain

**DOI:** 10.1101/2023.09.04.556176

**Authors:** Asimenia Gkogka, Susmita Malwade, Marja Koskuvi, Raj Bose, Sandra Ceccatelli, Jari Koistinaho, Jari Tiihonen, Martin Schalling, Samudyata, Carl M. Sellgren

## Abstract

Recent work in animal models suggests the role of oligodendrocyte precursor cells (OPCs) is not limited to oligodendrogenesis. Here, we report that OPCs derived from human induced pluripotent stem cells are capable of engulfing synaptosomes. We then develop a novel multi-lineage forebrain organoid in which OPCs alongside microglia display a spontaneous internalization of synaptic material in their phagolysosomes. Collectively, these findings suggest that OPCs contribute to synapse elimination in the developing human forebrain and provide a versatile platform for evaluating the interplay between microglia and OPCs in remodeling neuronal circuits in health and disease.

## INTRODUCTION

Oligodendrocyte precursor cells (OPCs) give rise to oligodendrocytes (OLs) which produce insulating sheaths of myelin around neuronal axons to enable efficient transmission of electrical impulses. In humans, OPC generation begins in early second trimester and progresses in several spatiotemporal waves towards birth.^1–4^ OPCs then expand and differentiate into OLs that initiate a highly coordinated program of myelination^5,6^, while surplus myelin and OPCs undergo refinement through microglial phagocytosis.^7–9^ However, a significant proportion of OPCs do not differentiate into OLs^10^ and may have important roles in the brain beyond oligodendrogenesis.^11–15^ Similar to microglia, recent work suggests that OPCs contribute to sculpting of the developing visual system in zebrafish and mouse through engulfment of excess synapses.^16–18^ The ability of OPCs to engulf thalamocortical inputs then significantly decreases after microglia depletion, indicating a close coordination between these two different glial subtypes in refining synapses in the developing brain.^16^

Models based on human induced pluripotent stem cells (iPSCs) have confirmed the capacity of human-derived microglia to engulf synaptic structures and have also indicated abnormal microglial synaptic pruning in various neurodevelopmental and neurodegenerative disorders.^19,20^ While protocols have been developed to generate OPCs from human iPSCs^21–25^, their phagocytic abilities have not been investigated, making it uncertain to what extent OPC-mediated synaptic pruning is a feature of the human brain development. Further, three-dimensional (3D) brain organoid models containing maturing OL-lineage cells in the presence of yolk sac-derived microglia are still missing, thus limiting our understanding of how these cell types interact in sculpting the developing neuronal circuits in health and disease contexts.

To address these questions, we first generated human iPSC-derived OPCs and employed a scalable high-throughput live-imaging system to explore OPC-mediated synaptic engulfment. Similar to what we previously observed for microglia^19^, we observed a stable internalization of synaptosomes in the phagolysosomes of iPSC-derived OPCs. To closely mimic the complexity of the developing human brain, we then established a human multi-lineage organoid model harbouring OL-lineage cells that transition through various developmental stages, in proximity to developing neurons, astroglia and microglia. Further, analyses of transcriptional data revealed that organoid-grown OPCs expressed key phagocytosis-related genes, while high-resolution imaging revealed that they are able of spontaneous uptake of synaptic material *in situ*. This suggests that OPCs contribute to the refinement of neuronal networks in the developing human brain and provide novel approaches to model OL lineage development and key functions in a human context.

## RESULTS

### Human iPSC-derived OPCs engulf synaptosomes

To study if human OPCs are capable of phagocytosing synaptic structures, we first generated OPCs from human iPSC lines (*n* = 3) using a previously published protocol.^21^ Briefly, neural progenitor cells (NPCs) were generated from iPSCs through embryoid bodies and rosette formation (days *in vitro*, DIV 0-21), followed by NPC culture in OPC induction medium for additional 10 DIV. At the end of OPC specification (DIV 31), cells displayed a typical bipolar morphology and stained positive for canonical OPC markers such as PDGFRα and OLIG2 (Figure 1A). Next, we characterized OPC phagocytosis in a previously validated 2D model of synaptic pruning^19,26^ by adding nerve terminals isolated from iPSC-derived neurons (synaptosomes) labelled with a pH-sensitive dye (pHrodo) that fluoresces when localized in acidic environments following phagocytosis, such as the phagolysosomal compartment. Fluorescence intensity measured at regular intervals and normalized to cell numbers demonstrated a robust internalization of synaptosomes by OPCs (Figure 1B-C), although the increase in phagocytic index over time and per cell was less pronounced as compared to microglia^19,26^. Fixation followed by immunostaining also confirmed an overlap of remaining pHrodo with the post-synaptic marker (PSD-95) and a marker for OPC lineage (PDGFRα) (Figure 1D-E).

**Figure 1.**
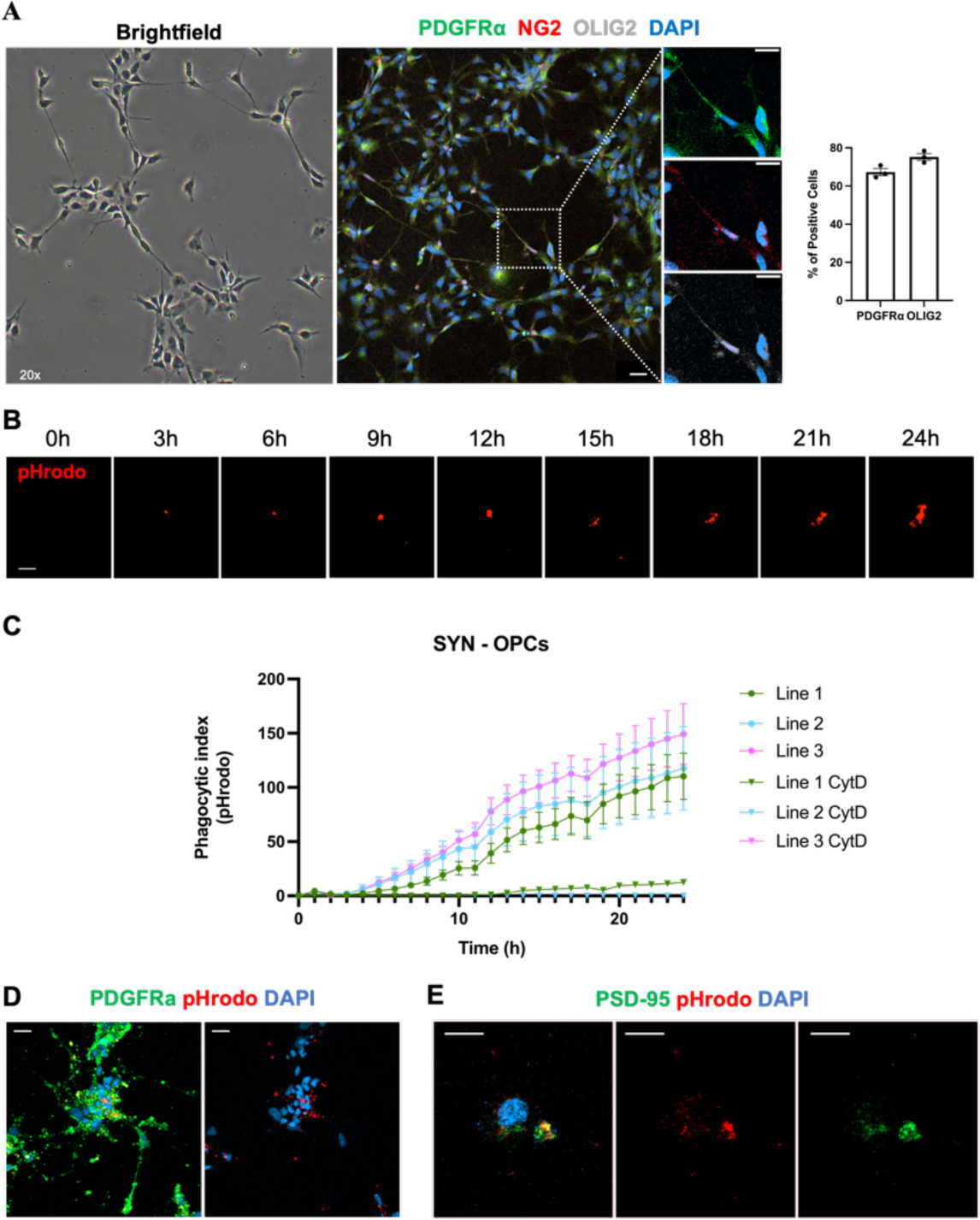
Human iPSC-derived OPCs internalize synaptosomes in the phagolysosome compartment. (A) Brightfield and fluorescent images of OPCs displaying typical bipolar OPC morphology and a high yield (> 65%) of PDGFRα+ and OLIG2+ cells in 2D culture. Data shown as mean ± s.e.m, *n* = 3 subjects. (B) Representative images from IncuCyte live imaging following uptake and accumulation of synaptosomes (SYN) in the phagolysosomes, as indicated by red fluorescence emitted from pHrodo, in an OPC. (C) Quantification of pHrodo-labeled SYN uptake by OPCs from cytochalasin D (10uM)-treated and untreated wells (*n* = 3 subjects). Phagocytic index represents the mean pHrodo+ area per OPC, data shown as mean ± s.e.m, across five images (20x) acquired every 1h per well. (D) Representative confocal images showing pHrodo-conjugated SYNs engulfed by PDGFRα+ OPCs (E) Representative confocal images showing co-localization of pHrodo and PSD-95+ puncta. Scale bars: 20 μm (A, B, D, E).

### Oligodendrogenesis in forebrain organoids with innate generation of microglia

Given that human iPSC-derived OPCs were capable of phagocytosing synaptosomes, we sought to recapitulate oligodendrogenesis in a relevant model of the developing human forebrain that would allow for OPC interactions with developing neuronal circuits as well as microglia. To this end, we combined iPSC-derived NPCs and primitive yolk sac macrophage progenitor cells (PMPs),^27^ capable of giving rise to microglia,^28^ and cultured the resulting 3D structures (DIV 0) with factors promoting OPC proliferation and differentiation, as well as factors needed for neuronal and microglial differentiation and survival (*n* = 3 subjects) (Figure 2A). Consistent with ganglionic eminence (GE) patterning in the presence of Sonic hedgehog (Shh) pathway agonists, we observed ventral progenitors (NKX2-1, OLIG2) ^29,30^ and the emergence of native OPC populations (PDGFRα)^31^ at 43 DIV and in the presence of developing microglia (IBA1) (Figure 2B). At 130 DIV, we observed post-mitotic neurons (MAP2), of both GABAergic (GABA) and glutamatergic (vGLUT1) identity, along with astroglia (GFAP, AQP4) (Figure 2C). From 43 DIV to 130 DIV, a substantial fraction of the OPCs had robustly differentiated into OLs as shown by the gradual decrease in PDGFRα+ precursor cells, from 24.0851% to 13.6455% of total cells at DIV 43 and 130, respectively (mean with ± s.e.m) and the simultaneous increase in the density of MBP+ OLs, from 5.99003 to 26.904 MBP+ μm^3^ per nuclei at DIV 43 and 130, respectively (mean with ± s.e.m) (Figure 2D). Further, OL processes were also found in proximity to axons wrapped in myelin (Figure 2E).

**Figure 2.**
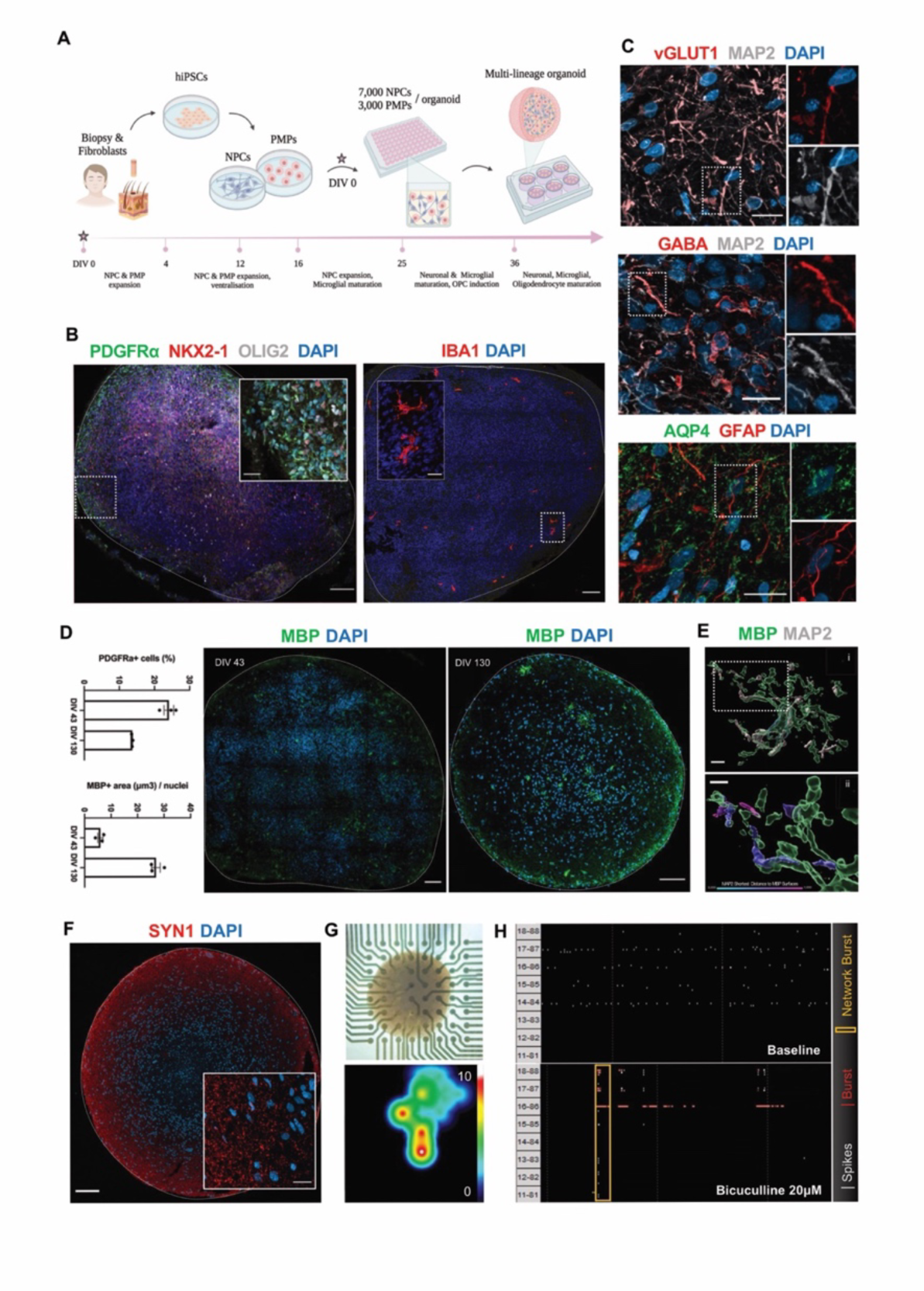
Generation and characterization of multi-lineage organoids containing OL lineage cells and microglia. (A) Graphical scheme summarizing the protocol. (B) Representative immunohistochemistry (IHC) images at 43 DIV showing ventral progenitors (NKX2-1, OLIG2) and OPCs (PDGFRα), in the presence of innately developing microglia (IBA1). (C) Representative IHC images at 130 DIV displaying mature neurons (MAP2) of glutamatergic (vGLUT1) (top) and GABAergic (GABA) (middle) identity along with astroglia (GFAP, AQP4) (bottom). (D) IHC images of whole organoid sections and bar plots showing % of PDGFRα+ OPCs and MBP+ area, indicating myelination, normalized with the total number of nuclei (DAPI) at 43 and 130 DIV. Data shown as mean ± s.e.m. (E) (i) Imaris-based volumetric reconstruction of an MBP+ cell wrapping MAP2+ neuronal processes with myelin, (ii) zoomed region showing statistically color-coded distance (cyan to magenta) between MAP2 and MBP structures with blue indicating surface overlap. (F) Representative IHC image of pre-synaptic elements (SYN1) across a whole organoid slice at 120 DIV. (G) Bright field image of an intact multilineage organoid mounted on a 64-electrode MEA plate at 100 DIV (top) and representative activity heat map showing spike rate (spike/s) (bottom). (H) Representative raster plots of organoids in a 50s timeframe at baseline (top) and after dosing with a GABAA receptor antagonist, bicuculline (20μM) (bottom). Scale bars: 100 μm (B, C, D), 10 μm (C, Ei), 20 μm (B, F) and 4 μm (Eii).

Next, to assess synaptogenesis in our organoids we stained for the pre-synaptic marker Synapsin 1 (SYN1) and observed abundant synapses organized along the organoid periphery (Figure 2F). We then recorded electrical activity of 100+ DIV organoids using multielectrode arrays (MEA) and observed spontaneous firing, indicating neuronal activity (Figure 2G). Further, blockade of GABAergic transmission in the presence of bicuculline^32^ increased the spontaneous neuronal firing and synchronized bursting (Figure 2H), then demonstrating integration of excitatory and inhibitory neurons into functional neuronal networks within the organoids.

### Characterization of multi-lineage forebrain organoids

To further profile cellular diversity in the generated multi-lineage forebrain organoids, we performed large-scale droplet-based single nucleus RNA sequencing on freshly dissociated organoids at 250 DIV (4 organoids per line, *n* = 2 subjects), and captured high-quality transcriptomic profiles of 10,645 nuclei (3624 median genes). Briefly, nuclei from individual subjects were demultiplexed and integrated followed by dimensionality reduction and unsupervised graph-based clustering resulting in 20 distinct cellular clusters (Figure 3A, S1A-D). Reference mapping of obtained clusters with previously characterized datasets of primary human fetal brain and organoid models, combined with inspection of top differentially expressed genes (Table S1) and canonical cell type signatures (Figure 3B) aided in identifying key neurodevelopmental cell types including neurons (*STMN2, RBFOX3*), OL-lineage cells (*PDGFR*α*, OLIG2, OLIG1*), microglia (*AIF1, C1QA*), ependymal cells (*FOXJ1, PIFO*), as well as radial glia populations that could be resolved into cycling (*TOP2A*, *MKI67*), non-cycling (*FOXG1*, *HOPX, GLI3)* and glioblast progenitors (*EGFR*, *TNC*, *GFAP*) (Figure S1E,F). Transcriptomic correlations to previously published primary human fetal brain datasets^33,34^ revealed regional similarities to the developing telencephalon of the second-trimester fetal brain (Figure S2A-C). Comparison to other brain region-specific organoid models^35^ revealed transcriptional overlap primarily with the forebrain-directed organoid model (Figure S2D), consistent with our protocol design. Importantly, microglia developed within the organoid model showed high transcriptional similarity with primary fetal microglia from the developing forebrain and exhibited a core microglia signature (Figure 3B, S2E). Next, to evaluate identities of the neuronal clusters, we utilized a reference dataset spanning interneuron specification in the developing primate GEs^36^ to score eminence-specificity which indicated the presence of neuronal lineages arising from medial (M), lateral (L) and caudal (C) GE progenitors (Figure S2F). Accordingly, multiple clusters of glioblasts expressed previously identified regional markers such as *MEIS2* (LGE-derived), *NR2F1* and *NR2F2* (CGE-derived), but not *NKX2-1* (MGE-derived, detectable at 43 DIV but not 250 DIV; Figure 2B) at the profiled developmental timepoint, then recapitulating the temporal dynamics of GE neurogenesis (Figure S2G).^37,38^ Similarly, differentiated neuronal clusters expressed markers corresponding to newborn interneurons (*DLX1*, *DLX2*, *DLX6, GAD1, GAD2, BCL11B*), including parvalbumin (PV) interneuron precursors (*ST18*, *ETV1*)^36,39^ (Figure S2G) and other interneuron subtypes (*SST*, *NPY*, *RELN*, *CALB2*) (Figure S2H).

**Figure 3.**
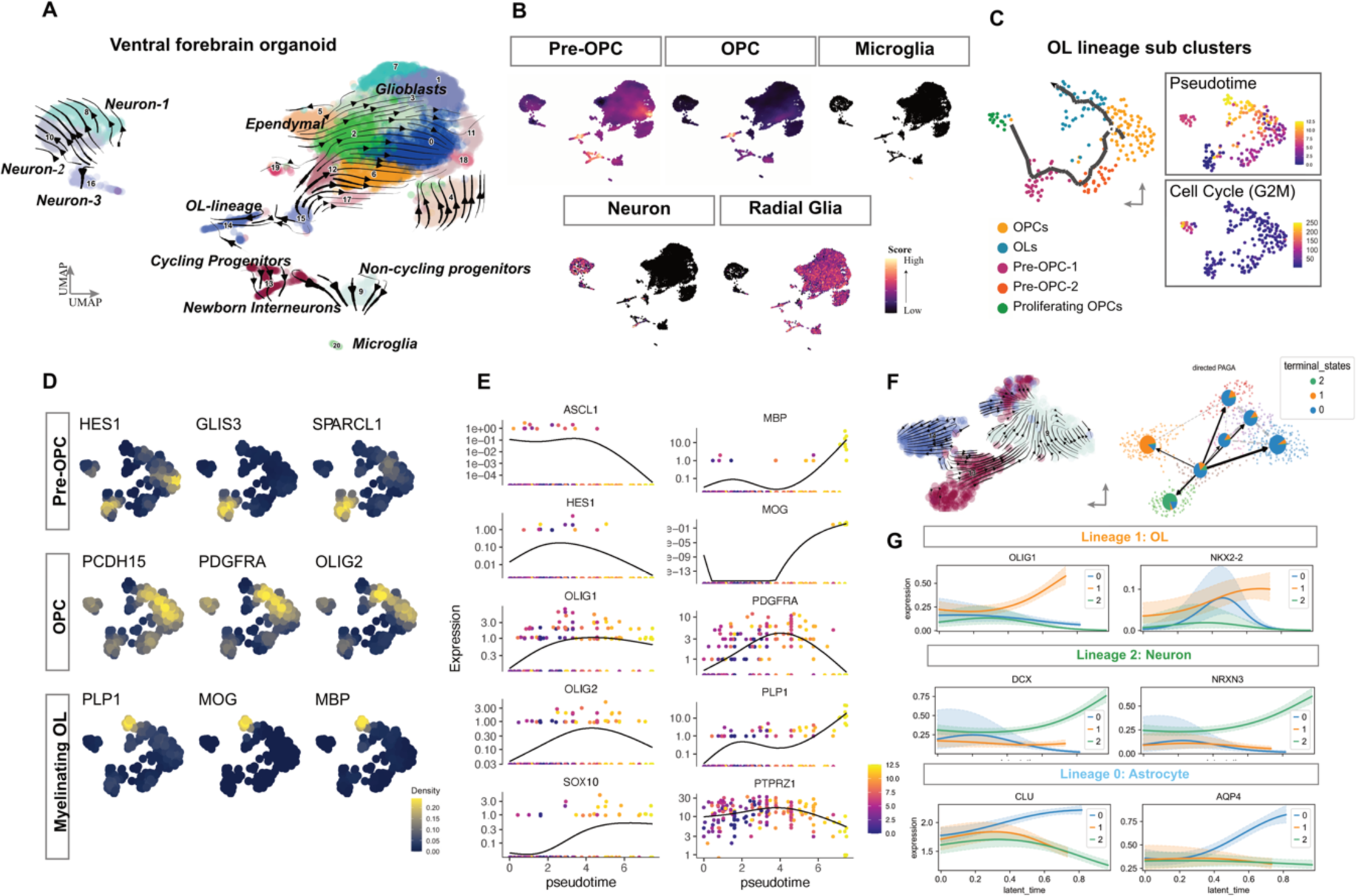
Deep characterization of multi-lineage forebrain organoids using single-cell transcriptomics. (A) UMAP plot depicting 10,645 nuclei transcriptomes from forebrain organoids (DIV 250) across 20 clusters. Arrows denote RNA velocities as streamlines derived from dynamical modeling of the organoid projected on the UMAP embedding. Numbers denote the cluster IDs. Each dot represents a cell colored according to its cluster ID. Quality control for the clustering is displayed in Figure S1. Computed marker genes specific to each cluster ID are provided in Table S1. (B) UMAP plots displaying the distribution of joint density of expression of genes representing cell type-specific signatures for Pre-OPC and OPC (van Bruggen et al^3^), Microglia (Patir et al^42^), Radial Glia *(HES1, HES6, SPARCL1, SOX6, DLX1, DLX5, EBF2, EOMES, NR2F2)*. The color bar represents joint kernel density. (C) UMAP plot (left) representing the 5 subclusters obtained from cluster 14 (OL-lineage) in Figure 3A. Each dot represents a cell colored by the respective subcluster. UMAP plots (right) of the cells colored by computed pseudotime and the cell cycle score for G2M phase. Black line represented the estimated trajectory and arrow represents the end of pseudotime. (D) UMAP plots displaying the expression of sub-state specific markers of the OL lineage to identify the subclusters. Color scale represents expression kernel density. (E) Expression of canonical OL lineage markers in individual cells ordered by pseudotime, denoted on the x-axis. Black lines indicate fitted spline curves. (F) UMAP embedding (top) displaying the dynamical modeling of isolated cluster 14, 13 and 9. Cells confidently assigned to each terminal macrostate identified using CellRank are highlighted according to the legend (bottom) on a directed partition-based graph abstraction with pie charts indicating aggregated fate probabilities. (G) Trajectory-specific gene expression trends across latent time obtained by fitting generalized additive models according to fate probabilities for each terminal macrostate, as seen in 3F.

We then thoroughly characterized the OL-lineage cells to study lineage-specific dynamics within the organoids. We assigned an OL-lineage score to all cells and selected cluster 14 for displaying the highest score (Figure 3C). Upon sub clustering, we then obtained five transcriptomic states along the differentiation spectrum, corresponding to pre-OPCs (*HES1*, *GLIS3, SPARCL1*), cycling (*MKI67*) and non-cycling OPCs (*BCAN*, *PDGFR*α), as well as OLs consisting of myelinating cells (*MBP, MOG, PLP1*) (Figure 3D-E). The identified OPC to OL states largely overlapped with O4+ cells from ventral forebrain organoids^40^ as well as primary OL cells from the fetal forebrain^3,34,41^ (Figure S3A, B). Further, dynamic modeling using RNA velocity highlighted OL lineage-specific genes across the OPC trajectory and revealed an additional cluster (13) associated to OPCs (Figure S3C). We identified cluster 13 to consist of multipotent progenitors as they showed high probabilistic fates towards astroglia, newborn interneurons and OL lineage with a pre-OPC-like signature (Figure 3F, S3C). We computed terminal states of this subset of progenitors and identified putative driver genes of the three potential fates using Markov state modelling and confirmed lineage-specific markers (Figure 3F). In an unsupervised approach, we then confirmed the multi-lineage potential of this progenitor subset with *LRRC4C*, *OLIG1* and *PCDH15* identified as the top driver genes towards the OL lineage, *DLX6-AS1, DCX* and *NRXN3* driving the neuronal fate, and *CLU, ADGRV1, NEAT1* and *AQP4* driving the progenitors towards the astroglial terminal state (Figure S3D).

### Cellular crosstalk patterns in multi-lineage forebrain organoids

By taking advantage of spatio-temporal development of different cell types in the generated organoids, we utilized hierarchical clustering of the transcriptomic data to determine global outgoing and incoming communication across major signaling pathways. The resulting patterns recapitulated functions of broad cell types such as neurons (pattern #2), microglia (pattern #3), OL lineage and newborn interneurons (pattern #4) as well as progenitors (pattern #1) (Figure S3E, F). Top contributors to outgoing signaling pattern of OL lineage cells were involved in cell adhesion/migration (ITGB2, CDH1, SEMA4)^43,44^ and immune regulation (TNF, CLEC, MHC-II),^11,15,45^ whereas main signaling receivers in the OL lineage were involved in trophic support (PDGF, PSAP),^46,47^ myelination (MPZ), and synaptic adhesion (NGL, NCAM, NEGR).^48^ For microglia both outgoing and incoming pathways were related to glial survival (CSF, TGFB, EGF)^49–51^ and immune surveillance (COMPLEMENT, IL1, CD226, APP, MIF),^52–55^ while neurons sent and received signals involving migration and positioning (RELN, L1CAM, SEMA5, SEMA6, EPHA),^56^ self-tolerance (THBS)^57^ and synaptic assembly (NLG, NRXN, NGL, NECTIN),^48^ and exclusively received signaling related to synaptic maintenance (COMPLEMENT).^54^ In sum, this supported that the model recapitulated cellular crosstalk patterns critical for neurodevelopmental processes.

### Spontaneous internalization of synapses by OPCs in multi-lineage forebrain organoids

We then analyzed how microglia and OPCs interacted with neuronal circuits in organoids after 130 DIV. Similar to microglia-containing undirected whole brain organoids,^58^ high-resolution confocal imaging followed by 3D volumetric reconstruction revealed uptake of post-synaptic terminals (GEPH) by microglia (IBA1), including overlap with the phagolysosomal marker LAMP2 (Figure 4Α). Next, we focused on OPCs and observed that PDGFRα+ cells made extensive contacts with synapses and displayed synaptic material in various stages of engulfment, from intact synaptic units consisting of both pre- and post-synapses (SYN1+, GEPH+) to complete internalization of GEPH+ terminals within LAMP2+ compartments (Figure 4B, C). Interestingly, analyses of synaptic uptake revealed both OPCs and microglia spontaneously internalized similar volumes of post-synaptic terminals (GEPH) normalized to their cellular volumes (0.00464466 and 0.00538254 total GEPH volume per cell volume internalized by OPCs and microglia, respectively, mean ± s.e.m) (Figure 4D). To compare the phagocytic machinery of OPCs and microglia, we then analyzed expression profiles of genes related to phagocytic and lysosomal function^18^ (Figure 4E). Both microglia and OPCs within the organoids expressed genes known to be involved in microglial phagocytosis (*PTPRJ*, *DOCK2*, *CD302, ITGAV),*^59–62^ as well as astrocyte-mediated synaptic refinement (*MERTK*, *ABCA1*),^63,64^ and certain phagocytosis-promoting genes (*LRP1*, *MEGF10, RAP1GAP*).^63,65–67^ Genes supporting exposure to eat-me signals (*XK*-related),^68^ were notably more highly expressed in OPCs (Figure 4E). Taken together, these results indicate that human OPCs harbor phagocytic machinery and that the model is suitable for studies aiming to precisely defining the molecular mechanisms that governs the synapse engulfment of human OPCs.

**Figure 4.**
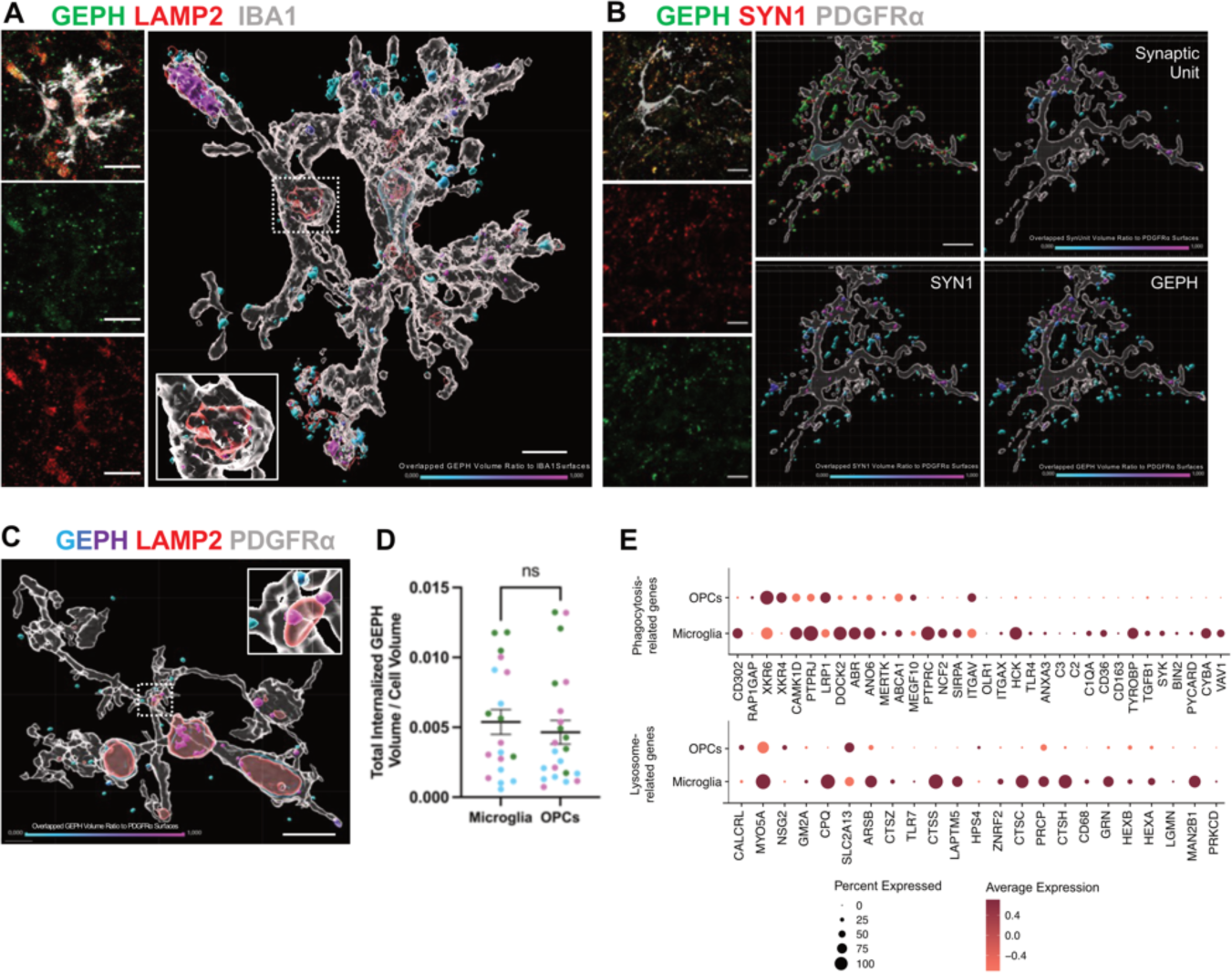
Spontaneous synaptic engulfment by different glial cells, including OPCs, within multilineage forebrain organoids. Imaris-based volumetric reconstruction of (A) post-synaptic inhibitory (GEPH) objects within microglial cells (IBA1), some of which are localized within phagolysosomes (LAMP2), and (B) pre-synaptic (SYN1) and post-synaptic (GEPH) terminals as well as intact synaptic units (when the shortest distance between SYN1 and GEPH is lower than 0.2 μm) within OPCs (PDGFRα), (C) some of which are localized within phagolysosomes (LAMP2). Statistically color-coded bar indicating overlapped volume ratio with PDGFRα+ surface (magenta = complete internalization). (D) Quantification of total internalized volume of GEPH+ objects, which display at least 90% volumetric overlap with the cell surface, normalized with total cell volume for at least 6 OPCs and microglia across three subject lines (*n* = 3). (E) Dot plots of phagolysosome-related genes expressed by OPCs and microglia. Individual data points are shown with bars representing mean ± s.e.m, two-tailed Mann–Whitney *U* test. Scale bars: 5 μm (A), 10 μm (B).

## DISCUSSION

Here, we first demonstrate that OPCs generated from human iPSCs are capable of engulfing isolated synaptic structures and provide a scalable high-throughput live-imaging system for quantification of OPC-mediated synaptic engulfment. We then develop and validate a novel multi-lineage human forebrain organoid model that includes microglia and multiple stages of OL lineage development and confirm that OPCs are capable of spontaneous internalization of synapses from functional neuronal circuits.

Selective removal of excess synapses is an essential neurodevelopmental process that contributes to the refinement of neuronal circuits.^69^ Microglia utilize complement signalling to identify redundant synapses, and defects in this recognition system has been observed in neurodevelopmental as well as in neurodegenerative disorders.^70^ Our findings reveal a novel role for human OPCs in eliminating synapses and raises questions about the proportion of OPCs engaged in this process at any developmental time point and the phagocytic machinery these cells rely on, as well as its potential implications for injury and disease. By allowing for integration of microglia, through addition of PMPs, the organoid model also supports physiologically relevant process of coordinated synapse removal between OPCs and microglia,^16^ while still making it is possible to distinguish cell-specific contribution. Thus the model systems used in this study provide opportunities for application to a variety of brain disorders while also being compatible with mechanistic studies.

### Limitations of the study

While 3D cultures more closely capture the *in vivo* cytoarchitecture as compared to 2D cultures, the cellular states of versatile cell types such as microglia are difficult to completely capture in an *in vitro* context. Recently, microglia-containing organoids were successfully transplanted into mouse brain with improved cellular characteristics.^71^ The advantages of this platform are likely to pertain also to organoids with a complete glial cell type repertoire. In line with previous single cell characterizations of brain organoids, high-resolution RNA sequencing (250 DIV) indicated correspondence to roughly second trimester. While further *in vitro* culture and/or transplantation is likely to support successive maturation, it is important to note that the brain organoid platform at this stage most accurately captures the developing rather than the mature human brain.

## Methods

Collection of biopsies, generation, and maintenance of human iPSCs Dermal biopsies were collected under standardized conditions (as approved by the Regional Ethical Review Boards in Stockholm, Sweden) and fibroblast cultures were established as previously described^19^ and tested negative for mycoplasma. Fibroblasts were reprogrammed into iPSCs using either mRNA reprogramming as previously described (1 line)^19^ or viral reprogramming using the CytoTune-iPS 2.0 Sendai Reprogramming Kit, according the manufacturer’s instructions (3 lines). iPSC lines (two females and two males) were expanded in mTeSR1 on Matrigel-coated plates and purified via magnetic cell sorting (MACS) with Anti-TRA-1-60 MicroBeads. At least three lines were used for all experiments.

### Generation and maintenance of PMPs

Human PMPs were generated from iPSCs in a Myb-independent manner, as previously reported,^27,28^ to recapitulate primitive hematopoiesis of tissue-resident macrophages, such as microglia, from yolk sac progenitors before the emergence of hematopoietic stem cells.^72^ Briefly, 10,000 iPSCs were cultured for 4 days into a V-bottom ultra-low attachment 96-well plate in pre-PMP embryoid body (EB) medium, which consisted of mTESR1 supplemented with bone morphogenetic protein 4 (BMP-4, 50 ng/ml), stem cell factor (SCF, 20 ng/ml), vascular endothelial growth factor (VEGF, 50 ng/ml) and Y-27632 rho-kinase inhibitor (10 μM). The resulting Ebs were transferred into tissue-culture treated 6-well plates containing PMP medium, which consisted of X-VIVO15 supplemented with interleukin-3 (IL-3, 25ng/ml), macrophage colony-stimulating factor (M-CSF, 100ng/ml), penicillin-streptomycin (P/S, 1X), β-mercaptoethanol (100 μΜ). Four weeks later, PMPs appeared in suspension, which were harvested for organoid generation.

### Generation and maintenance of NPCs

Human NPCs were generated from iPSCs, as previously described with minor modifications.^27^ Briefly, 4,000 iPSCs per well were cultured for 1 week into a V-bottom ultra-low-attachment 96-well plate to aggregate into embryoid bodies (EBs) using EB medium, consisting of Advanced DMEM/F12, N-2 (1X), B27 without vitamin A (1X), dorsomorphin (1 μΜ) and SB431542 (5 μM). Resulting EBs were plated on Matrigel-coated plates in medium composed of Advanced DMEM/F12, N-2 (1X) and laminin 521 (1 μg/ml) and developed in the form of rosettes for 1 week. Rosettes were then manually isolated from surrounding cells and expanded for 1 additional week in NPC medium, consisting of a 1:1 mixture of Neurobasal and Advanced DMEM/F12, supplemented with N-2 (1X), B27 without vitamin A (1X), basic fibroblast growth factor (bFGF, 20 ng/ml), human leukemia inhibitory factor (hLIF, 10 ng/ml), CHIR99021 (3 μM), SB431542 (2 μM) and Y-27632 rho-kinase inhibitor (10 μM).

### Generation and maintenance of OPCs

Human OPCs were generated from iPSCs, as previously described^21^ with minor modifications. Briefly, following differentiation from iPSCs, NPCs were expanded for one week in Neural expansion medium, composed of a 1:1 mixture of Neurobasal and Advanced DMEM/F-12, N2 (1X) and B27 without vitamin A (1X), supplemented with bFGF (20 ng/ml) and epidermal growth factor (EGF, 20 ng/ml). Using Anti-A2B5 MicroBeads, A2B5+ cells were then isolated and plated on a fibronectin (25 μg/ml)-coated dish and cultured in OPC-like medium, composed of a 1:1 mixture of Neurobasal and Advanced DMEM/F12, supplemented with N-2 (1X), B27 without vitamin A (1X), platelet-derived growth factor (PDGF-AA, 10 ng/ml), insulin-like growth factor 1 (IGF-1, 10 ng/ml), Sonic hedgehog (SHH, 100 ng/ml), forskolin (5μM), N-acetyl cysteine (60 μg/ml), bFGF (20 ng/ml) and Noggin (20 ng/ml) for 10-15 days.

### Immunocytochemistry

Cells were fixed in 4% paraformaldehyde (PFA) for 15 min at RT, washed twice with phosphate buffered saline (PBS). Permeabilization and blocking was performed for 30 min at room temperature (RT) in 3% bovine serum albumin (BSA), 0.1% Triton X-100 (VWR) in PBS. Cells were washed twice in PBS and incubated overnight at 4°C with antibody solution composed of 3% BSA, 0.1% Triton X-100 in PBS and the desired primary antibodies. Subsequently, cells were washed twice in PBS and incubated with antibody solution and appropriate secondary antibodies for 1 hour at RT in the dark. Stained samples were washed twice in PBS, incubated with DAPI (1:500) for 5 min and mounted with Fluorescence Mounting Medium. Confocal laser-scanning microscopes Zeiss LSM800, LSM900-Airy, with ZEN2009 software were used for imaging. ImageJ was used for image processing and analysis.

### Purification and labelling of SYNs

SYNs were isolated from NPC-derived neuronal cultures using Syn-PER Synaptic Protein Extraction Reagent, as described previously in Sellgren *et al*., 2017. Briefly, neurons were lifted off the plate with the use of a cell scraper and the sample was centrifuged at 1,200 g for 10 min. Supernatant was collected and centrifuged at 15,000 g for 20 min, and SYNs were collected in the form of a pellet and resuspended in medium containing 10% DMSO to protect against freezing artifacts. For real-time live-cell assays, SYNs were thawed, resuspended in 0.1 M sodium carbonate buffer (pH 9.0) and labeled according to the manufacturer’s instructions with an amine-reactive and pH-sensitive dye (pHrodo red). Labelled SYNs were sonicated for 20 min to avoid clumping and resuspended in sodium carbonate buffer before its addition to the cells.

### Synaptic vesicle endocytosis by 2D OPCs

The IncuCyte ZOOM live imaging system was used at 37 °C and 5% CO2, as previously described. Briefly, sonicated and pHrodo-labeled SYNs were added to OPCs seeded at a 13,000 cells per well density in 96-well imaging plates and five images per well were acquired with the 10x objective every 1 hour for a total period of 24 hours (3 technical replicates, 5 images per replicate for each line). Images were pre-processed with the IncuCyte ZOOM software (version 2016A) and exported as PNG files. The phagocytic index refers to the ratio of the total red fluorescence emitted per well divided by the total number of OPCs present in each well every hour. Live imaging was followed by fixation of cells with 4% PFA for 15 min at RT followed by washing twice with PBS for immunocytochemistry assays and confocal microscopy.

### Generation and maintenance of multi-lineage organoids

Organoids were generated by co-culturing 7,000 NPCs and 3,000 PMPs derived from three iPSC lines per organoid, as described previously^27^ with some modifications. The resulting EBs were maintained into a V-bottom ultra-low-attachment 96-well plate in a 1:1 mixture of NPC and PMP medium for the first 4 DIV. PMP medium was composed of X-VIVO 15, supplemented with IL-3 (25 ng/ml) and M-CSF (100 ng/ml). For the next 11 days (DIV 5-15), organoids were cultured with a mixture of 1:1 PMP and pre-OPC medium, and on DIV 8 transferred to ultra-low-attachment 6-well plates and kept on the orbital shaker at a speed of 80 rpm. Pre-OPC medium consisted of basal medium, supplemented with bFGF (20 ng/ml) and EGF (20 ng/ml). Basal medium comprised of 1:1 mixture of Neurobasal and Advanced DMEM/F12, GlutaMax (1X), MEM Non-Essential Amino Acids (1X), human insulin (25 μg/ml), β-mercaptoethanol (100 μM), P/S (1X), N-2 supplement (1X), and B-27 supplement without vitamin A (1X). During DIV 16-24, organoids were cultured in pre-OPC medium, supplemented with microglial maturation factors interleukin-34 (IL-34, 100 ng/ml) and granulocyte-macrophage colony-stimulating factor (GM-CSF, 10 ng/ml). Further, to pattern these organoids towards the ventral forebrain fate, Wnt pathway inhibitor IWP-2 (5μΜ) was added to the medium during DIV 4-24 and the small molecule smoothened agonist (SAG, 1μM) during DIV 12-24. From DIV 25 until DIV 35, they were cultured in OPC medium, consisting of basal medium with triiodo-L-thyronine (T3, 60 ng/ml), biotin (100 ng/ml), neurotrophin-3 (NT-3, 20 ng/ml), cyclic AMP (cAMP, 1μM), hepatocyte growth factor (HGF, 5 ng/ml), IGF-1 (10 ng/ml), PDGF-AA (10 ng/ml), supplemented with IL-34 (100 ng/ml), GM-CSF (10 ng/ml) along with the neuronal maturation factors brain-derived neurotrophic factor (BDNF, 20 ng/ml) and glial cell-derived neurotrophic factor (GDNF, 20 ng/ml). From DIV 36 onwards they were grown in OL medium consisting of 2:1:1 mixture of BrainPhys neuronal medium, Neurobasal and Advanced DMEM/F12, respectively, with GlutaMax (1X), MEM (1X), human insulin (25 μg/ml), β-mercaptoethanol (100 μM), P/S (1X), N-2 (1X) and B-27 without vitamin A (1X), supplemented with T3 (60 ng/ml), biotin (100 ng/ml), cAMP (1μM), ascorbic acid (20 μg/ml), IL-34 (100 ng/ml), GM-CSF (10 ng/ml), BDNF (20 ng/ml) and GDNF (20 ng/ml). Media changes were performed daily until DIV 15, every other day until DIV 43 and every 3 days from DIV 44 onwards. Minimum 10 organoids were generated per line.

### Cryopreservation

Following fixation in 4% PFA for 45 min at RT, the organoids were washed twice in PBS for 5 min and transferred to 30% sucrose solution (w/vol, Sigma) for overnight incubation at 4 °C. Upon removal of sucrose, organoids were embedded in optimal cutting temperature (OCT) compound and stored at −80 °C until used for cryosectioning. Organoid blocks were then allowed to equilibrate to sectioning temperature in the cryostat chamber for 30 min prior to sectioning, when 18-μm-thick sections of frozen organoid tissue were prepared using a Leica cryostat and stored at −20 °C until use. The optimal temperature of the blade and the chamber was at −20°C and multiple serial sections were collected from each organoid, allowing for exploration of several markers across different regions of the organoids.

### Immunohistochemistry

Cryosections were washed with PBS, followed by permeabilization, and blocking in 3% BSA, 0.1% Triton X-100 in PBS for 1 hour at RT. The sections were then washed twice in PBS and incubated overnight at 4°C with primary antibodies diluted in blocking solution. Upon washing twice with PBS, the cryosections were incubated in a mixture of appropriate secondary antibodies diluted in blocking solution for 2 hours at RT in the dark. Stained samples were washed twice with PBS, incubated with DAPI for 5 min and mounted on glass slides using Fluorescence Mounting Medium. For anti-PDGFRα, an initial permeabilization step with 0.2% Triton X-100 in PBS for 10 min was added before blocking, a quenching step with TrueBlack® Plus Lipofuscin Autofluorescence Quencher diluted in 70% ethanol for 30sec was added before mounting. Imaging was performed on confocal laser-scanning microscope Zeiss LSM800 using ZEN2009 software. ImageJ was used for image processing and CellProfiler was used for cellular quantifications. For myelination related quantifications in Figure 2D, the total MBP+ surface (μm^3^) was normalized to the total number of nuclei (DAPI) in 1-μm z-sections of whole organoid slices from *n* = 3 cell lines. For data presentation, the volumetric reconstruction of an MBP+ cell was statistically color-coded, ranging from cyan = 0 to magenta = 1, using the “shortest distance to surface” function with the MAP2+ neuronal processes.

### Spontaneous engulfment quantification within multi-lineage organoids

40x images of DIV 130 organoid sections stained for OPCs and/or microglia were pre-processed using ImageJ, by applying the “Enhance contrast” command so that 0.1% of pixels would be saturated, and the “Mean filter” with pixel radius of 1.5 µm. Then, each cell of interest was cropped into its own image stack file, which was used in downstream analysis. In Imaris 10.0 (BitPlane, UK), volumetric reconstructions of the fluorescence images were created using the “Surfaces” objects to reconstruct the cell (PDGFRα, IBA1), the synaptic elements (GEPH, SYN1, PSD-95) and the lysosomes (LAMP2) of interest using the fluorescence as a reference. The synaptic units were defined as the pre-synaptic element surfaces (SYN1) with lower than 0.2 μm shortest distance from the post-synaptic element surfaces (GEPH) by using the filtering command and by creating a new surface consisting of pre- and post-synaptic elements within this distance range. The synaptic particles were defined as internalized when ≥ 90% of their volume ratio overlapped with the defined cell surface. The volumes of the cell and the internalized surfaces were collected from the statistics tab in Imaris. The total internalized volume was then normalized to a cell’s volume to represent the amount of engulfment by each cell. For data presentation, the synaptic elements and unit surfaces were statistically color-coded, ranging from cyan = 0 to magenta = 1, based on the overlapped volume ratio with the cell surface, indicating internalization.

### Multi-electrode array (MEA) recording

6-well MEA plates (Axion Biosystems) were coated with 0.1% Polyethyleneimine solution for an hour, washed 2x with water, and one organoid per well was placed on the electrode grid in 100ul of media supplemented with laminin 521 (10 μg/ml) for a couple of hours to facilitate attachment. For the next 48h, the organoid maturation medium was supplemented with laminin 521 (1 μg/ml), followed by media changes every 2-3 days. The recordings were performed using a Maestro MEA system with AxIS Software Spontaneous Neural Configuration (Axion Biosystems). Spikes were detected with AxIS software using an adaptive threshold crossing set to 5.5 times the standard deviation of the estimated noise for each electrode. Pharmacological interventions were performed on organoids plated on 6-well MEA plates using bicuculline (20 μM), with baseline recordings obtained immediately before and after 15 minutes of adding each compound.

### Dissociation of organoids and snRNA-seq

Four organoids per subject line (*n* = 2) were processed for single nuclei (sn) RNA sequencing at DIV 250. To isolate single nuclei from fresh organoid tissue samples, organoids were subjected to cold mechanical dissociation with glass pipettes to obtain a single-cell suspension. Dissociated cells were centrifuged at 500 g for 5 minutes at 4°C. The cell pellet was washed and exposed to a freshly prepared lysis buffer (as demonstrated in protocol CG000366 RevD, 10x Genomics) followed by centrifugation at 500g for 5 minutes at 4°C. To remove cellular debris, the nuclei suspension was further subjected to an iodoxinol gradient (Optiprep; 25% and 29%) with centrifugation at 13500 g for 20 minutes at 4°C. The resulting nuclei pellet was resuspended in diluted nuclei buffer and observed under a Leica brightfield microscope at 40X magnification to determine nuclei quality. The nuclei concentration was manually counted using trypan blue and a Neubauer counting chamber. Isolated nuclei were equally pooled and loaded onto the Chromium controller (10x Genomics) with a target recovery of 10000 cells. The assay was processed according to the Chromium Next GEM Single Cell 3’ Reagent Kits v3.1(Dual Index) user guide (RevD). Upon GEM generation, barcoding, cleanup and cDNA amplification, a single barcoded library was constructed and sequenced on Illumina NovaSeq S1-100 (v1.5, 2 x 100 bp).

### snRNA-seq data processing and analyses

Raw sequencing data obtained as fastq files were processed using the cellranger (v6.1.2) pipeline with read alignment to the human GRCh38/hg38 reference genome. Barcodes and sorted bam files generated by cellranger were used for genotype demultiplexing of the two individual cell lines by souporcell package, with k=2 and a common variants file (limited to >=2% MAF) of GRCh38. Droplets identified as empty or containing ambient RNA were excluded from further analysis. Each cell was assigned to the respective individual cell line. UMI count matrices were processed in R using the Seurat package. All cells were further subjected to quality control metrics according to the number of transcripts (0000) and genes (>200 and <11000) captured, the percentage of mitochondrial transcripts (5%). Genes expressed in less than 3 cells were excluded. Cells were scored for the cell cycle phase in Seurat and the difference between the S phase and G2M phase scores was calculated. Doublets were identified and filtered out using DoubletFinder (pN=0.25, pK=0.22). Stressed cells within the organoid were identified using the Gruffi package. Post quality control, gene expression per cell was normalized and scaled using SCTransform function (3000 features) to account for technical artifacts and the cell cycle difference was regressed. Batch effects were evaluated and cells originating from individual cell lines were normalized individually and integrated in Seurat using FindIntegrationAnchors and IntergateData functions. To reduce dimensionality, UMAP algorithm was applied to the integrated dataset with 30 dimensions in PCA space as input. Unsupervised graph-based clustering was performed in Seurat using K-nearest neighbors’ algorithm and Louvain algorithm (at 1.0 and 1.4 resolution) to find unique cellular clusters.

### RNA velocity and cellular trajectories

BAM files generated by cellranger were used as input to Velocyto package to obtain the spliced and unspliced counts matrices for all genes. RNA velocity analysis was performed using the scVelo package in Python with pre-computed dimensionality reduction and clustering. Using the integrated data, the first order and second-order moments were computed (default settings) and the velocities were calculated using the likelihood-based dynamical model. The velocity graph was visualized on the UMAP embedding as streams. Using the dynamical model, cellular trajectories were inferred along with their initial and terminal states within the dataset using CellRank toolkit in Python. Fate probabilities for each cell and driver genes for the OL-lineage were computed using tr.lineage drivers function.

### Differential expression and cell type annotations

Differential gene expression across identified clusters was performed using FindAllMarkers function in Seurat using the MAST test (MALAT1 and mitochondrial genes were excluded). Multiple-testing correction was performed using the Benjamini-Hochberg method. The cluster markers along with known canonical marker gene expressions were visualized on the UMAP embeddings. Annotations were finalized based on a combination of unsupervised markers, transcriptomic correlations, and post-integration label transfers from multiple publicly available snRNA-seq datasets. External datasets from Braun *et al*. (https://doi.org/10.1101/2022.10.24.513487) Cameron *et al*. (10.1016/j.biopsych.2022.06.033), Nowakowski *et al*. (2017), Marton *et al*., (10.1038/s41593-018-0316-9), and Amin *et al*. (https://doi.org/10.1101/2023.05.31.541819) were directly downloaded either as pre-processed R objects or gene expression count matrices with available annotations from respective publications. For cluster comparisons, clusters were transcriptomically correlated (Pearsons’s correlation) to external reference datasets to aid cell type annotations using the clustifyr package. Further, the dataset from this study was integrated using the Harmony package with external organoid datasets used as a reference to perform classification and transfer cluster labels using TransferData function onto the query data.

### Cellular crosstalk within organoids

Cell-cell communication networks across identified cellular clusters were computed and visualized using the CellChat toolkit (v1.6.0). A CellChat object was created from the Seurat object with discrete clusters and signaling genes. The ligand-receptor pairs for humans were extracted from CellChatDB database. Overexpressed ligand-receptor interactions were identified and the ‘trimeans’ method was used to calculate average gene expression per cell group. The communication probability of all ligand-receptor interactions linked to known signaling pathways from the database was computed and the aggregated weighted-directed network across cell types of interest was visualized as circle plots and chord diagrams. To identify global communication patterns (incoming and outgoing), unsupervised hierarchical clustering (k=4) of signaling pathways across cell groups was performed. Significant interactions between cell types were determined using permutation testing and significance was considered at p<0.05.

### Data and code Availability

Single-cell analysis and visualization were performed in R (v 4.2.2) and Python (v3.7.12) on macOS (Ventura13.4). Raw and processed data files will be made available at the GEO database. The code reproducing the single-cell analysis is available at https://github.com/orgs/SellgrenLab/oligodendrocyte-organoid. For other data analysis, GraphPad Prism software was used.

## ACKNOWLEDGMENT

We thank the study participants and acknowledge the technical support provided by Biomedicum Imaging Core (BIC) facility at Karolinska Institutet. The computations/data handling was enabled by resources provided by provided by the National Academic Infrastructure for Supercomputing in Sweden (NAISS) and the Swedish National Infrastructure for Computing (SNIC) at UPPMAX, Uppsala University. We are also grateful to Julschen Majkowitz for valuable feedback on the manuscript. This work was generously supported by grants from Erling Persson Foundation (C.M.S), Hjärnfonden (C.M.S.: FO2022-0135), the regional agreement on medical training and clinical research between Stockholm County Council (C.M.S.: 2017-02559), and Karolinska Institutet (C.M.S.: KID).

## AUTHOR CONTRIBUTIONS

S and C.M.S conceived the study. S, A.G generated organoids with help from M.K and performed all the experiments. S, S.M performed dissociations and prepared libraries for sequencing with help from A.G. S.M analysed the transcriptomic data. MEA experiments were done in collaboration with R.B and S.C. J.K and J.T provided some of the IPSC lines. S, A.G, S.M and C.M.S interpreted the data and wrote the manuscript with input from other co-authors.

