## Supplementary Figure S1 for "Oligodendrocyte precursor cells engulf synapses in a model of the developing human forebrain"

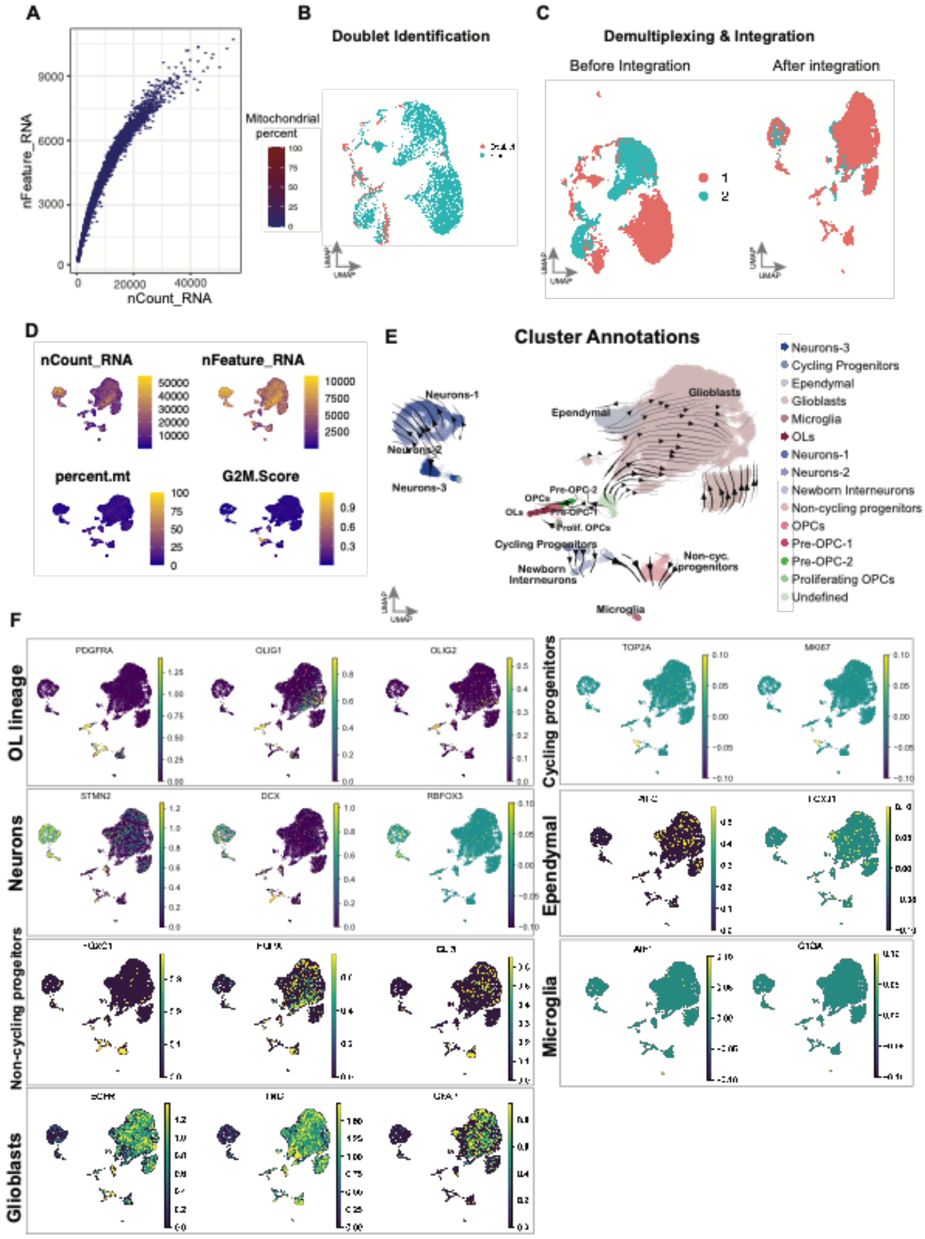


**Figure S1.** **(A)** Scatter plot denoting number of transcripts captured (x-axis) versus the number of genes captured (y-axis) per cell within the dataset after quality control. Data points represent individual cells colored by the percentage of mitochondrial transcripts. **(B)** UMAP plot showing each cell colored by the doublet status assigned by DoubletFinder. Assigned doublets were filtered out for downstream applications. **(C)** UMAP embedding of the dataset with each cell colored by subject line, demultiplexed by genotype (see methods), as a confounder (left). Cells from individual cell lines were integrated (see methods) to obtain the final embedding for downstream analysis (right). **(D)** UMAP plots of the final embedding showing quality control metrics such as number of transcripts (nCount_RNA), genes (nFeature_RNA), percentage of mitochondrial transcripts (percent.mt), and G2M phase score for individual cells, showing uniform distribution and no confounding effects. **(E)** UMAP plot visualizing the distribution of clusters colored by the cell type annotations used for downstream analyses. Black lines and arrows denoted RNA velocity streamlines. **(F)** UMAP plots showing the distribution of canonical marker gene expression for broad cell types during brain development across individual cells. Color scale represents the average scaled expression for single genes.
