## Supplementary Figure S2 for "Oligodendrocyte precursor cells engulf synapses in a model of the developing human forebrain"

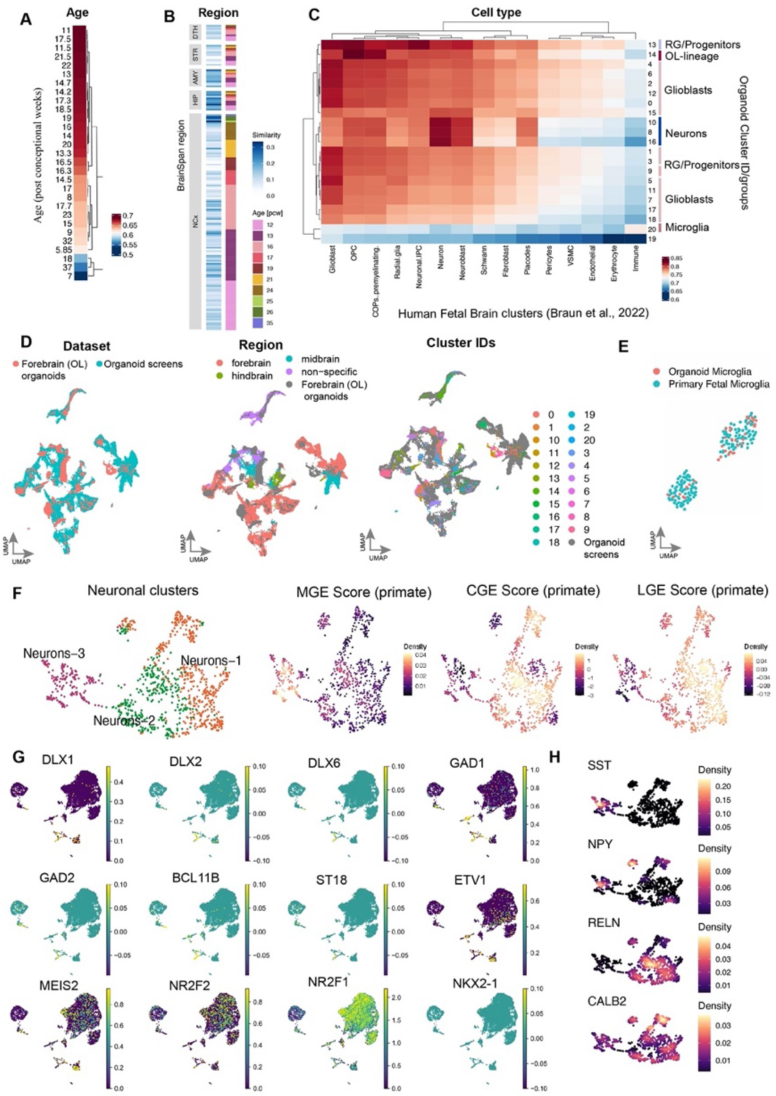


**Figure S2.** Reference mapping utilising spearman-ranked correlation of forebrain organoids to reference transcriptomes of **(A)** primary fetal cortex and GE (Nowakowski et al) for fetal age, **(B)** spatial similarity to human micro dissected brain from BrainSpan (https://www.brainspan.org/), and **(C)** large-scale dataset of human fetal brain (Braun et al[33]) for cell type similarities. **(D)** UMAP embedding of the integrated space of forebrain organoid dataset (this study) with region-directed brain organoids dataset (Amin et al), including forebrain, midbrain and hindbrain organoids. UMAP plots show cells colored by dataset (left), regional specification (middle), and cluster IDs from figure 3A (right). Legends for each plot are provided on the bottom right respectively. **(E)** UMAP plot of the integrated space of microglia (cluster 20) from forebrain organoids with human fetal microglia (Nowakowski et al) showing an overlap across 2 microglia states from the fetal brain. **(F)** UMAP embedding of re-clustered neuronal cell subset (comprising of cluster IDs: 8,10,16 from figure 3A) followed by visualizing the cells scored according to the GE-specific signatures obtained from the developing macaque brain (Zhao et al). Joint density estimation of the scores is visualized on the UMAP embedding of the neuronal subset. UMAP plots visualize the **(G)** average scaled expression of known GE region-specific marker genes across all cells, and **(H)** the expression density of key neuronal subtype marker genes across the neuronal clusters shown above in S2F.
