## Supplementary Figure S3 for "Oligodendrocyte precursor cells engulf synapses in a model of the developing human forebrain"

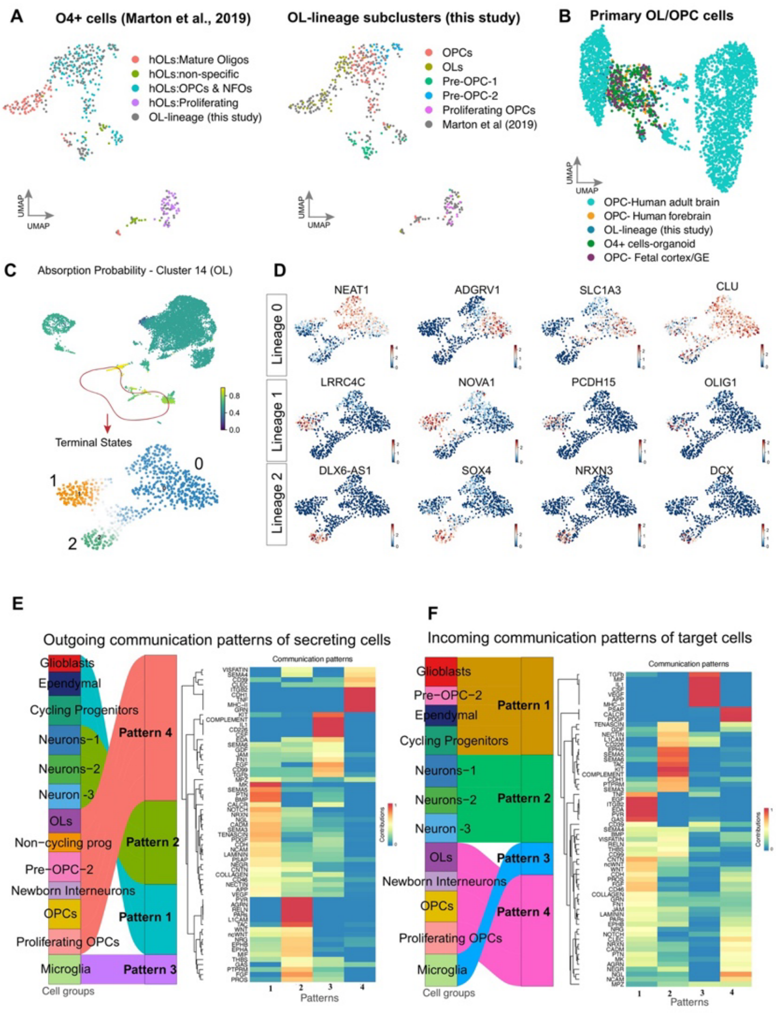


**Figure S3.** **(A)** UMAP plot of the integrated space of OL-lineage cells (cluster 14) from forebrain organoids (this study) with a reference dataset of organoid-isolated O4+ve cells (Marton et al). Cells are colored by the original annotations of the reference dataset (left) and the annotated OL sub-clusters (figure 3C) from this study (right). **(B)** UMAP plot of the integrated space of single-cell datasets of OPCs from human adult brain (Hodge et al[42]), human forebrain (van Bruggen et al[3]), human developing cortex (Nowakowski et al[32]), organoid-isolated O4+ve cells (Marton et al[39]), and OL-lineage (cluster 14) cells from forebrain organoids from this study. **(C)** UMAP embedding visualizing absorption probabilities of individuals cells, estimating the likelihood of transitioning to terminal macrostate of cluster 14 (figure 3A) representing the OL cluster, for the whole dataset. Cells highlighted by the red circle (cluster 9,13,14) were isolated (seen in figure 3F) for computing fates, resulting in 3 terminal macrostates identified by Cellrank (bottom). 30 most confidently assigned terminal state cells are highlighted by colors per predicted lineage (bottom). **(D)** Top 4 lineage driver genes identified for each lineage visualized on the UMAP embedding from figure 3F. Color scale indicated computed correlations between fate probabilities and gene expression. Inferred latent communication patterns of signaling pathways used by cell groups **(E)** secreting (outgoing) to other groups as well as **(F)** received (incoming) from other groups within the organoids visualized as alluvial plots. The patterns are shown as pattern #1 (P1), pattern #2 (P2), pattern #3 (P3), pattern #4 (P4). Significant signaling pathways contributing to each pattern are displayed as a heatmap respectively. Scale bar indicates communication probability scores. Significance was tested at $p<0.05$.
