## Supplementary Table S1 for "Oligodendrocyte precursor cells engulf synapses in a model of the developing human forebrain"

| p_val | avg_log2FC | pct.1 | pct.2 | p_val_adj | cluster | gene |
|---|---|---|---|---|---|---|
| 4,635E-246 | 1,07060037 | 0,943 | 0,781 | 1,379E-241 | 0 | TRPM3 |
| 4,691E-282 | 1,05704434 | 0,951 | 0,711 | 1,396E-277 | 0 | SLC1A2 |
| 7,432E-212 | 0,91501682 | 0,868 | 0,551 | 2,212E-207 | 0 | SLC4A4 |
| 1,749E-206 | 0,72033039 | 0,743 | 0,368 | 5,205E-202 | 0 | SLC6A11 |
| 1,153E-159 | 0,70801461 | 0,935 | 0,742 | 3,433E-155 | 0 | RYR3 |
| 2,783E-204 | 0,70141114 | 0,909 | 0,635 | 8,284E-200 | 0 | C1orf61 |
| 1,889E-202 | 0,68293493 | 0,974 | 0,8 | 5,622E-198 | 0 | SPARCL1 |
| 4,082E-118 | 0,65937752 | 0,758 | 0,476 | 1,215E-113 | 0 | ANOS1 |
| 6,472E-137 | 0,65258804 | 0,903 | 0,726 | 1,926E-132 | 0 | RGS7 |
| 2,896E-254 | 0,65200688 | 0,999 | 0,992 | 8,618E-250 | 0 | DTNA |
| 0 | 2,32377225 | 0,999 | 0,662 | 0 | 1 | DPP10 |
| 0 | 1,32312975 | 0,869 | 0,21 | 0 | 1 | DPP10-AS3 |
| 0 | 1,13784638 | 0,761 | 0,153 | 0 | 1 | AC093610.1 |
| 2,663E-248 | 1,09443026 | 0,963 | 0,698 | 7,925E-244 | 1 | HPSE2 |
| 0 | 0,94824006 | 0,646 | 0,131 | 0 | 1 | DPP10-AS1 |
| 2,357E-123 | 0,81653215 | 0,913 | 0,698 | 7,015E-119 | 1 | OBI1-AS1 |
| 8,462E-151 | 0,77564048 | 0,745 | 0,395 | 2,518E-146 | 1 | AC098656.1 |
| 3,746E-194 | 0,66692438 | 0,999 | 0,858 | 1,115E-189 | 1 | LINC01088 |
| 6,859E-181 | 0,65916685 | 0,993 | 0,871 | 2,041E-176 | 1 | CNTN1 |
| 1,87E-186 | 0,65165598 | 0,992 | 0,793 | 5,566E-182 | 1 | NAA11 |
| 1,0081E-87 | 0,45125819 | 0,997 | 0,89 | 3,0005E-83 | 2 | GLIS3 |
| 1,892E-102 | 0,43002497 | 0,998 | 0,861 | 5,6298E-98 | 2 | LINC01088 |
| 6,3529E-67 | 0,41927808 | 0,943 | 0,801 | 1,8908E-62 | 2 | OGFRL1 |
| 4,3446E-48 | 0,41132207 | 0,693 | 0,542 | 1,2931E-43 | 2 | PTCH2 |
| 8,2221E-88 | 0,40067869 | 0,978 | 0,798 | 2,4471E-83 | 2 | NAA11 |
| 6,4754E-33 | 0,39508407 | 0,761 | 0,618 | 1,9273E-28 | 2 | PTCHD4 |
| 3,0977E-51 | 0,37091319 | 0,988 | 0,942 | 9,2197E-47 | 2 | DGKI |
| 3,85E-40 | 0,35983935 | 0,854 | 0,709 | 1,1459E-35 | 2 | C8orf34 |
| 1,002E-69 | 0,34869276 | 0,978 | 0,846 | 2,9823E-65 | 2 | RGMA |
| 1,8107E-68 | 0,347855 | 0,988 | 0,869 | 5,3891E-64 | 2 | CALN1 |
| 1,169E-163 | 1,1138844 | 0,895 | 0,666 | 3,478E-159 | 3 | SLC7A14-AS1 |
| 2,854E-127 | 0,80354094 | 0,739 | 0,404 | 8,495E-123 | 3 | AC098656.1 |
| 6,755E-132 | 0,77814954 | 0,941 | 0,706 | 2,011E-127 | 3 | HPSE2 |
| 1,844E-100 | 0,74431838 | 0,531 | 0,266 | 5,4873E-96 | 3 | AC026316.5 |
| 2,168E-189 | 0,74029328 | 1 | 0,861 | 6,454E-185 | 3 | LINC01088 |
| 6,9662E-72 | 0,71578716 | 0,886 | 0,707 | 2,0734E-67 | 3 | OBI1-AS1 |
| 1,765E-171 | 0,70114101 | 0,993 | 0,797 | 5,255E-167 | 3 | NAA11 |
| 4,9525E-86 | 0,64259963 | 0,693 | 0,428 | 1,474E-81 | 3 | TNR |
| 1,0938E-60 | 0,60377685 | 0,598 | 0,385 | 3,2555E-56 | 3 | KCNU1 |
| 1,3684E-77 | 0,58720088 | 0,856 | 0,652 | 4,0729E-73 | 3 | AC093535.1 |
| 0 | 2,74062046 | 0,972 | 0,296 | 0 | 4 | IGFBP5 |
| 0 | 2,45832822 | 0,902 | 0,29 | 0 | 4 | VEGFA |
| 0 | 1,58472377 | 0,68 | 0,077 | 0 | 4 | HILPDA |
| 0 | 1,37384252 | 0,782 | 0,221 | 0 | 4 | ADM |
| 1,495E-196 | 1,31656411 | 0,714 | 0,343 | 4,45E-192 | 4 | PURPL |
| 1,113E-252 | 1,26297523 | 0,748 | 0,489 | 3,313E-248 | 4 | GBE1 |
| 0 | 1,15767837 | 0,902 | 0,557 | 0 | 4 | P4HA1 |
| 3,329E-199 | 1,09765966 | 0,759 | 0,347 | 9,907E-195 | 4 | LINC00836 |
| 0 | 1,08605241 | 0,997 | 0,969 | 0 | 4 | WSB1 |

| | | | | | | |
|---|---|---|---|---|---|---|
| 0 | 1,07789876 | 0,614 | 0,088 | 0 | 4 | PDK1 |
| 7,645E-273 | 2,22142329 | 0,531 | 0,12 | 2,275E-268 | 5 | GRIN2A |
| 0 | 2,02704879 | 0,894 | 0,202 | 0 | 5 | TTC6 |
| 0 | 1,84693131 | 0,773 | 0,335 | 0 | 5 | CFAP54 |
| 0 | 1,84344854 | 0,68 | 0,276 | 0 | 5 | AL139815.1 |
| 3,234E-210 | 1,67085324 | 0,889 | 0,625 | 9,624E-206 | 5 | AQP4-AS1 |
| 2,032E-211 | 1,48420301 | 0,549 | 0,103 | 6,049E-207 | 5 | FOXP2 |
| 4,154E-229 | 1,43952861 | 0,5 | 0,058 | 1,236E-224 | 5 | ARMC3 |
| 1,604E-224 | 1,41293574 | 0,969 | 0,711 | 4,775E-220 | 5 | C8orf34 |
| 1,306E-200 | 1,35586358 | 0,742 | 0,201 | 3,886E-196 | 5 | SLIT2 |
| 2,135E-240 | 1,25496359 | 0,634 | 0,23 | 6,356E-236 | 5 | CFAP43 |
| 1,446E-183 | 2,09079177 | 0,672 | 0,222 | 4,304E-179 | 6 | LHFPL3 |
| 1,212E-259 | 1,8593527 | 0,634 | 0,102 | 3,607E-255 | 6 | CATSPERB |
| 3,604E-141 | 1,74107238 | 0,812 | 0,636 | 1,073E-136 | 6 | FRMD5 |
| 1,596E-118 | 1,40928891 | 0,522 | 0,173 | 4,751E-114 | 6 | CHI3L1 |
| 5,513E-243 | 1,39033056 | 0,962 | 0,748 | 1,641E-238 | 6 | GRIA1 |
| 2,364E-111 | 1,28564056 | 0,979 | 0,84 | 7,037E-107 | 6 | DEC1 |
| 8,032E-149 | 1,25171795 | 0,91 | 0,579 | 2,391E-144 | 6 | SLC4A4 |
| 5,926E-199 | 1,2465278 | 0,626 | 0,131 | 1,764E-194 | 6 | TC2N |
| 6,658E-135 | 1,23749041 | 0,484 | 0,082 | 1,982E-130 | 6 | LHFPL3-AS1 |
| 1,714E-132 | 1,1867245 | 0,893 | 0,555 | 5,103E-128 | 6 | SHISA9 |
| 2,414E-252 | 1,87748748 | 0,993 | 0,509 | 7,186E-248 | 7 | ROBO2 |
| 1,418E-181 | 1,8219047 | 0,978 | 0,695 | 4,219E-177 | 7 | DPP10 |
| 6,309E-144 | 1,54500895 | 0,98 | 0,716 | 1,878E-139 | 7 | OBI1-AS1 |
| 1,863E-151 | 1,20086107 | 0,941 | 0,42 | 5,544E-147 | 7 | AC098656.1 |
| 4,4462E-96 | 1,16779837 | 0,517 | 0,159 | 1,3233E-91 | 7 | PI15 |
| 6,4652E-86 | 1,14670599 | 0,465 | 0,144 | 1,9242E-81 | 7 | AC011632.1 |
| 4,179E-126 | 1,1062988 | 0,786 | 0,277 | 1,244E-121 | 7 | DPP10-AS3 |
| 2,019E-124 | 0,97333647 | 0,445 | 0,071 | 6,009E-120 | 7 | ADAMTSL3 |
| 1,4106E-93 | 0,92288068 | 0,897 | 0,518 | 4,1985E-89 | 7 | EFEMP1 |
| 2,2885E-96 | 0,9212913 | 0,616 | 0,175 | 6,8112E-92 | 7 | SLC14A1 |
| 0 | 4,74402608 | 0,998 | 0,227 | 0 | 8 | GALNTL6 |
| 0 | 4,02175995 | 1 | 0,221 | 0 | 8 | LRRTM4 |
| 0 | 3,93467717 | 0,991 | 0,187 | 0 | 8 | TPH2 |
| 0 | 3,64663202 | 0,97 | 0,18 | 0 | 8 | SGCZ |
| 0 | 3,45235234 | 0,998 | 0,553 | 0 | 8 | SYT1 |
| 0 | 3,43226623 | 1 | 0,303 | 0 | 8 | IQCJ-SCHIP1 |
| 0 | 3,40937256 | 0,998 | 0,134 | 0 | 8 | CNTNAP5 |
| 0 | 3,37559477 | 0,975 | 0,153 | 0 | 8 | ADARB2 |
| 0 | 3,35897576 | 0,991 | 0,345 | 0 | 8 | GRIA4 |
| 0 | 3,24155149 | 0,998 | 0,27 | 0 | 8 | GRIK2 |
| 0 | 4,09187129 | 0,994 | 0,209 | 0 | 9 | ADGRV1 |
| 0 | 2,82367275 | 0,979 | 0,564 | 0 | 9 | AL589740.1 |
| 0 | 2,36376841 | 0,909 | 0,236 | 0 | 9 | LUCAT1 |
| 0 | 2,17388955 | 0,959 | 0,813 | 0 | 9 | PPP1R9A |
| 5,212E-111 | 2,16001401 | 0,691 | 0,217 | 1,551E-106 | 9 | NRG1 |
| 2,759E-283 | 1,97466105 | 0,826 | 0,219 | 8,213E-279 | 9 | LRRC3B |
| 0 | 1,89694094 | 0,974 | 0,504 | 0 | 9 | PTN |
| 3,325E-204 | 1,82365599 | 0,991 | 0,959 | 9,896E-200 | 9 | NRXN1 |
| 5,653E-260 | 1,79786159 | 0,735 | 0,057 | 1,683E-255 | 9 | ADAMTS3 |

| | | | | | | |
|---|---|---|---|---|---|---|
| 0 | 1,77608414 | 0,918 | 0,212 | 0 | 9 | LTBP1 |
| 0 | 3,55570114 | 0,991 | 0,31 | 0 | 10 | IQCJ-SCHIP1 |
| 0 | 3,26794426 | 0,964 | 0,099 | 0 | 10 | SLC35F3 |
| 2,789E-282 | 3,15869355 | 0,94 | 0,163 | 8,3E-278 | 10 | ADARB2 |
| 3,02E-296 | 3,15431011 | 0,94 | 0,197 | 8,989E-292 | 10 | TPH2 |
| 0 | 3,11313012 | 0,988 | 0,143 | 0 | 10 | CNTNAP5 |
| 8,535E-224 | 3,00620678 | 0,877 | 0,238 | 2,54E-219 | 10 | ROBO1 |
| 3,001E-261 | 3,00104739 | 0,871 | 0,205 | 8,931E-257 | 10 | RALYL |
| 1,414E-176 | 2,98661472 | 0,757 | 0,215 | 4,208E-172 | 10 | NRG1 |
| 2,881E-290 | 2,96232166 | 0,964 | 0,247 | 8,574E-286 | 10 | CNTNAP2 |
| 0 | 2,91634117 | 0,997 | 0,213 | 0 | 10 | ANK3 |
| 2,429E-119 | 1,64893147 | 0,639 | 0,365 | 7,229E-115 | 11 | ALK |
| 1,4863E-49 | 1,24125016 | 0,89 | 0,723 | 4,4237E-45 | 11 | OBI1-AS1 |
| 1,5757E-85 | 1,1862615 | 0,856 | 0,427 | 4,6898E-81 | 11 | GABBR2 |
| 3,984E-165 | 1,14271136 | 0,538 | 0,034 | 1,186E-160 | 11 | RSPO2 |
| 6,8157E-98 | 1,11980659 | 0,97 | 0,821 | 2,0286E-93 | 11 | SPARCL1 |
| 2,3475E-75 | 1,09036918 | 0,957 | 0,607 | 6,9868E-71 | 11 | LUZP2 |
| 4,3612E-56 | 0,91853962 | 0,799 | 0,375 | 1,298E-51 | 11 | IL1RAPL1 |
| 1,537E-134 | 0,8873396 | 0,428 | 0,016 | 4,574E-130 | 11 | AC025508.1 |
| 2,5633E-62 | 0,87794917 | 1 | 0,959 | 7,6292E-58 | 11 | NRXN1 |
| 7,2287E-32 | 0,86215442 | 0,896 | 0,742 | 2,1515E-27 | 11 | SLC1A2 |
| 2,696E-139 | 1,64684791 | 0,951 | 0,797 | 8,024E-135 | 12 | CADPS |
| 1,006E-121 | 1,21062553 | 0,58 | 0,063 | 2,994E-117 | 12 | NIBAN1 |
| 3,9607E-73 | 1,09809746 | 0,259 | 0,038 | 1,1788E-68 | 12 | AL357507.1 |
| 7,2732E-94 | 1,09396019 | 0,967 | 0,821 | 2,1647E-89 | 12 | PSAP |
| 1,3112E-28 | 1,07598421 | 0,63 | 0,435 | 3,9025E-24 | 12 | RBFOX1 |
| 1,9265E-77 | 0,98232642 | 0,642 | 0,375 | 5,7338E-73 | 12 | DMGDH |
| 2,9488E-35 | 0,90995358 | 0,745 | 0,541 | 8,7765E-31 | 12 | FTL |
| 3,2259E-51 | 0,90152096 | 0,449 | 0,22 | 9,6011E-47 | 12 | ROR1 |
| 1,6312E-51 | 0,89998902 | 0,753 | 0,554 | 4,8549E-47 | 12 | SQSTM1 |
| 1,5628E-69 | 0,86648145 | 0,765 | 0,422 | 4,6515E-65 | 12 | IARS |
| 9,285E-259 | 3,51726431 | 0,799 | 0,396 | 2,763E-254 | 13 | NRXN3 |
| 2,429E-156 | 2,83412962 | 0,456 | 0,011 | 7,229E-152 | 13 | DLX6-AS1 |
| 4,407E-209 | 2,50448315 | 0,657 | 0,1 | 1,312E-204 | 13 | DNAH11 |
| 1,163E-208 | 2,43178989 | 0,979 | 0,854 | 3,461E-204 | 13 | SOX2-OT |
| 1,649E-166 | 2,39964684 | 0,64 | 0,111 | 4,909E-162 | 13 | AC068308.1 |
| 1,392E-149 | 1,93217062 | 0,77 | 0,31 | 4,144E-145 | 13 | KCNH7 |
| 0 | 1,92714413 | 0,766 | 0,173 | 6,487E-308 | 13 | SOX4 |
| 0 | 1,90521729 | 0,82 | 0,161 | 0 | 13 | SYNE2 |
| 4,199E-152 | 1,83862718 | 0,356 | 0,081 | 1,25E-147 | 13 | AC007402.1 |
| 9,609E-182 | 1,82538818 | 0,686 | 0,032 | 2,86E-177 | 13 | PDGFRA |
| 1,68E-301 | 3,81933291 | 0,995 | 0,233 | 5E-297 | 14 | LHFPL3 |
| 1,841E-257 | 3,46952775 | 0,964 | 0,166 | 5,479E-253 | 14 | PCDH15 |
| 1,758E-170 | 2,99188931 | 0,959 | 0,249 | 5,232E-166 | 14 | OPCML |
| 3,268E-211 | 2,98282913 | 0,927 | 0,091 | 9,728E-207 | 14 | AL353784.1 |
| 3,888E-237 | 2,76388308 | 0,912 | 0,149 | 1,157E-232 | 14 | CHRM3 |
| 3,348E-259 | 2,72096929 | 0,99 | 0,793 | 9,965E-255 | 14 | KCND2 |
| 6,636E-289 | 2,56192566 | 0,995 | 0,854 | 1,975E-284 | 14 | SOX2-OT |
| 3,682E-205 | 2,53150978 | 0,917 | 0,089 | 1,096E-200 | 14 | LHFPL3-AS1 |
| 4,074E-168 | 2,4161335 | 0,793 | 0,111 | 1,212E-163 | 14 | AC068308.1 |

| | | | | | | |
|---|---|---|---|---|---|---|
| 1,304E-258 | 2,40388089 | 0,959 | 0,49 | 3,88E-254 | 14 | MMP16 |
| 1,969E-215 | 2,75916781 | 0,427 | 0,196 | 5,862E-211 | 15 | PLCG2 |
| 3,149E-141 | 1,83775958 | 0,302 | 0,102 | 9,373E-137 | 15 | AL627171.2 |
| 2,63E-154 | 1,39156791 | 0,578 | 0,471 | 7,828E-150 | 15 | TMSB4X |
| 5,128E-149 | 1,37103106 | 0,323 | 0,147 | 1,526E-144 | 15 | TMSB10 |
| 8,259E-171 | 1,33722898 | 0,427 | 0,416 | 2,458E-166 | 15 | HSP90AA1 |
| 7,928E-171 | 1,22202938 | 0,297 | 0,251 | 2,36E-166 | 15 | H3F3B |
| 1,7964E-78 | 1,17300353 | 0,635 | 0,496 | 5,3468E-74 | 15 | FTH1 |
| 5,5877E-63 | 1,1291685 | 0,703 | 0,543 | 1,6631E-58 | 15 | FTL |
| 1,882E-151 | 1,08859503 | 0,438 | 0,356 | 5,601E-147 | 15 | EEF1A1 |
| 7,27E-162 | 1,06665659 | 0,37 | 0,237 | 2,164E-157 | 15 | RPL13A |
| 0 | 3,88874673 | 0,973 | 0,422 | 0 | 16 | KCNIP4 |
| 6,705E-227 | 3,78166814 | 0,967 | 0,257 | 1,995E-222 | 16 | CNTNAP2 |
| 9,515E-188 | 3,3216393 | 0,956 | 0,246 | 2,832E-183 | 16 | ROBO1 |
| 1,962E-227 | 3,21784606 | 0,951 | 0,319 | 5,839E-223 | 16 | TENM2 |
| 8,817E-203 | 2,94029118 | 0,995 | 0,185 | 2,624E-198 | 16 | CCSER1 |
| 2,54E-157 | 2,91668492 | 0,868 | 0,189 | 7,56E-153 | 16 | CNTN5 |
| 3,595E-148 | 2,77626869 | 0,929 | 0,158 | 1,07E-143 | 16 | CDH18 |
| 3,528E-207 | 2,68792916 | 0,912 | 0,15 | 1,05E-202 | 16 | NKAIN2 |
| 5,016E-160 | 2,6649688 | 0,879 | 0,067 | 1,493E-155 | 16 | AC117461.1 |
| 9,417E-170 | 2,63812078 | 1 | 0,224 | 2,803E-165 | 16 | ANK3 |
| 6,066E-110 | 2,27948759 | 0,748 | 0,089 | 1,806E-105 | 17 | AC079352.1 |
| 1,8541E-41 | 2,02895062 | 0,785 | 0,242 | 5,5183E-37 | 17 | LHFPL3 |
| 2,1075E-48 | 1,53000989 | 0,991 | 0,593 | 6,2725E-44 | 17 | SLC4A4 |
| 4,4115E-62 | 1,43941299 | 0,916 | 0,173 | 1,313E-57 | 17 | PCDH15 |
| 7,6073E-53 | 1,37643448 | 1 | 0,929 | 2,2642E-48 | 17 | NOVA1 |
| 2,6091E-40 | 1,35551353 | 0,972 | 0,693 | 7,7656E-36 | 17 | GRID2 |
| 2,1808E-45 | 1,35509394 | 0,869 | 0,435 | 6,4909E-41 | 17 | GABBR2 |
| 1,176E-43 | 1,33168916 | 0,86 | 0,236 | 3,5002E-39 | 17 | NXPH1 |
| 4,141E-59 | 1,2874438 | 0,869 | 0,155 | 1,2325E-54 | 17 | CHRM3 |
| 1,3978E-28 | 1,27998446 | 0,533 | 0,1 | 4,1604E-24 | 17 | LHFPL3-AS1 |
| 8,584E-85 | 2,41602391 | 0,733 | 0,185 | 2,5548E-80 | 18 | AC002069.2 |
| 1,8911E-90 | 2,02128104 | 0,571 | 0,171 | 5,6284E-86 | 18 | AC003991.1 |
| 1,3027E-82 | 1,6357791 | 0,952 | 0,661 | 3,8771E-78 | 18 | EGFR |
| 1,67E-54 | 1,33630603 | 0,495 | 0,089 | 4,9704E-50 | 18 | STEAP4 |
| 2,1933E-42 | 1,13019694 | 0,895 | 0,697 | 6,5278E-38 | 18 | LYRM2 |
| 5,7142E-42 | 1,12638303 | 0,771 | 0,47 | 1,7007E-37 | 18 | AKNAD1 |
| 3,6334E-31 | 1,08857219 | 0,829 | 0,569 | 1,0814E-26 | 18 | SMOC1 |
| 1,3856E-30 | 1,07674228 | 0,762 | 0,414 | 4,1239E-26 | 18 | GLCCI1 |
| 3,9701E-30 | 1,06616245 | 0,971 | 0,834 | 1,1816E-25 | 18 | CST3 |
| 9,661E-22 | 1,01292039 | 0,733 | 0,292 | 2,8754E-17 | 18 | PAK3 |
| 9,9423E-62 | 2,67528207 | 0,9 | 0,633 | 2,9591E-57 | 19 | KCNMB2 |
| 1,5837E-47 | 2,61480382 | 0,65 | 0,153 | 4,7135E-43 | 19 | AL008633.1 |
| 1,3693E-83 | 2,56078347 | 0,7 | 0,113 | 4,0754E-79 | 19 | AP001977.1 |
| 3,4609E-47 | 2,19759195 | 0,95 | 0,594 | 1,0301E-42 | 19 | KCNMB2-AS1 |
| 8,4597E-69 | 1,98367994 | 0,65 | 0,06 | 2,5179E-64 | 19 | AC093607.1 |
| 6,975E-61 | 1,98224942 | 0,375 | 0,048 | 2,076E-56 | 19 | AL078602.1 |
| 1,0263E-30 | 1,97173566 | 0,825 | 0,332 | 3,0546E-26 | 19 | SGO1-AS1 |
| 3,9616E-51 | 1,9359625 | 0,825 | 0,386 | 1,1791E-46 | 19 | NUP210L |
| 5,1227E-42 | 1,87418708 | 0,875 | 0,645 | 1,5247E-37 | 19 | EYS |

| | | | | | | |
|---|---|---|---|---|---|---|
| 1,1202E-39 | 1,86083186 | 0,675 | 0,048 | 3,334E-35 | 19 | COL21A1 |
| 2,644E-188 | 4,40102224 | 1 | 0,023 | 7,868E-184 | 20 | FCGBP |
| 3,733E-136 | 4,12732951 | 1 | 0,014 | 1,111E-131 | 20 | CD74 |
| 0 | 4,05748363 | 0,972 | 0,103 | 0 | 20 | CCDC26 |
| 8,703E-113 | 4,03206835 | 0,917 | 0,005 | 2,59E-108 | 20 | CD207 |
| 2,94E-46 | 3,21855339 | 0,861 | 0,057 | 8,7503E-42 | 20 | GABRG3-AS1 |
| 1,475E-98 | 3,07957808 | 1 | 0,008 | 4,39E-94 | 20 | HLA-DRA |
| 3,095E-124 | 3,07120292 | 0,972 | 0,32 | 9,21E-120 | 20 | KCNQ1OT1 |
| 2,5464E-73 | 2,90913256 | 0,917 | 0,031 | 7,5787E-69 | 20 | EMCN |
| 1,2419E-95 | 2,88466442 | 0,972 | 0,004 | 3,6961E-91 | 20 | HLA-DRB1 |
| 4,672E-124 | 2,88007969 | 1 | 0,017 | 1,39E-119 | 20 | ATP8B4 |
